# Integration of Apocarotenoid Profile and Expression Pattern of *Carotenoid Cleavage Dioxygenases* during Mycorrhization in Rice

**DOI:** 10.1101/2023.02.24.529886

**Authors:** Cristina Votta, Jian You Wang, Nicola Cavallini, Francesco Savorani, Kit Xi Liew, Luisa Lanfranco, Salim Al-Babili, Valentina Fiorilli

**Author notes:** These authors contributed equally to this work (Cristina Votta, Jian You Wang).

## Abstract

Carotenoids are susceptible to degrading processes initiated by oxidative cleavage reactions mediated by Carotenoid Cleavage Dioxygenases that break their backbone, leading to products called apocarotenoids. These carotenoid-derived metabolites include the phytohormones abscisic acid and strigolactones, and different signaling molecules and growth regulators, which are utilized by plants to coordinate many aspects of their life. Several apocarotenoids have been recruited for the communication between plants and arbuscular mycorrhizal (AM) fungi and as regulators of the establishment of AM symbiosis. However, our knowledge on their biosynthetic pathways and the regulation of their pattern during AM symbiosis is still limited. In this study, we generated a qualitative and quantitative profile of apocarotenoids in roots and shoots of rice plants exposed to high/low phosphate concentrations, and upon AM symbiosis in a time course experiment covering different stages of growth and AM development. To get deeper insights in the biology of apocarotenoids during this plant-fungal symbiosis, we complemented the metabolic profiles by determining the expression pattern of *CCD* genes, taking advantage of chemometric tools. This analysis revealed the specific profiles of *CCD* genes and apocarotenoids across different stages of AM symbiosis and phosphate supply conditions, identifying novel markers at both local and systemic levels.

**Highlight:** Our study presents the profiles of *CCD* gene expression and apocarotenoids across different stages of AM symbiosis and Pi supply conditions and reveals novel AM markers at both local and systemic levels.

## Introduction

Carotenoids represent are a widespread class of tetraterpene (C_40_) lipophilic pigments, synthesized by all photosynthetic organisms, including bacteria, algae, and plants, and by numerous non-photosynthetic microorganisms (Moise *et al*., 2014; Nisar *et al*., 2015). In plants, carotenoids are essential constituents of the photosynthetic apparatus where they act as photo-protective pigments and take part in the light-harvesting process. Further, these pigments have ecological functions, providing flowers and fruits with specific colors and flavors that attract insects and other animals or act as a repellent for pathogens and pests (Cazzonelli, 2011).

The carotenoid structure, rich in electrons and conjugated double bonds, makes them susceptible to oxidation, which causes the breakage of their backbone and leads to a wide range of metabolites called apocarotenoids (Moreno *et al*., 2021). These compounds can be generated by non-enzymatic processes that are triggered by reactive oxygen species (ROS) (Harrison and Bugg, 2014; Ahrazem *et al*., 2016) or by the action of a ubiquitous family of non-heme iron enzymes, Carotenoid Cleavage Dioxygenases (CCDs) (Jia *et al*., 2018).

The genome of the model plant *Arabidopsis thaliana* encodes nine members of the CCD family, including five *NINE-CIS-EPOXY CAROTENOID CLEAVAGE DIOXYGENASES* (*NCED2, NCED3, NCED5, NCED6*, and *NCED9*) and four *CCD*s (*CCD1, CCD4, CCD7*, and *CCD8*) (Tan *et al*., 2003; Sui *et al*., 2013). In short, NCEDs catalyze the first step in abscisic acid (ABA, C_15_) biosynthesis, i.e. the cleavage of 9-*cis*-violaxanthin or 9’-*cis*-neoxanthin into the ABA precursor xanthoxin (Nambara and Marion-Poll, 2005; Ahrazem *et al*., 2016). CCD1 cleaves several carotenoids and apocarotenoids at different positions along their hydrocarbon backbone (Schwartz *et al*., 2001; Vogel *et al*., 2008; Ilg *et al*., 2009, 2014; Jia *et al*., 2018) generating volatiles, such as β-ionone and geranylacetone, and a diverse set of dialdehydes in fruits and flowers of many plant species (Moreno *et al*., 2021). CCD4 enzymes are known to produce apocarotenoid-derived pigments, flavors, and aromas *in planta*, but their cleavage specificities differ considerably from those of CCD1 enzymes (Schwartz *et al*., 2001; Auldridge *et al*., 2006; Ilg *et al*., 2009; McQuinn *et al*., 2015; Hou *et al*., 2016). In plants, two different forms of CCD4 are present (Huang *et al*., 2009; Mi and Al-Babili, 2019): the first one is common in *Citrus* and is involved in forming the pigment citraurin (3-hydroxy-β-apo-8′-carotenal, C_30_) formation by catalyzing a single cleavage reaction at the 7′,8′ double bond of zeaxanthin and β-cryptoxanthin (Ma *et al*., 2013; Rodrigo *et al*., 2013), while the other type cleaves bicyclic all-*trans*-carotenoids at the C9, C10 or C9’, C10’ double bond leading to apo-10’-carotenoids (C_27_) and the corresponding C_13_ cyclohexenones, e.g. β-ionone (Bruno *et al*., 2015, 2016). Recently, a Gardinia CCD4 enzyme (GjCCD4a) was shown to catalyze sequential cleavage of β-carotene and zeaxanthin at the C7, C8 and C8’, C9’ leading to crocetin dialdehyde, the precursor of the saffron pigment crocins, and to cyclocitral and 3-hydroxycyclocitral, respectively (Zheng et al., 2022).

The two CCD-subfamilies CCD7 and CCD8 are involved in the biosynthesis of the plant hormone strigolactones (SLs) (Wang *et al*., 2021). CCD7 cleaves 9-*cis*-β-carotene (C_40_), yielding β-ionone and 9-*cis*-β-apo-10’-carotenal (C_27_); while CCD8 converts 9-*cis*-apo-10’-carotenal (C_27_) *via* a combination of different reactions into the SL precursor carlactone (C_19_) and ω-OH-(4-CH3) heptanal (C_8_) (Alder *et al*., 2012; Chen *et al*., 2022). In addition, CCD7 may also catalyze the initial 9,10 cleavage required for mycorradicin synthesis (Floss *et al*., 2008).

A recent survey on plant genomes identified the Zaxinone Synthase (ZAS) as a representative for a further CCD subfamily, which is conserved in most land plants but missing in non-mycorrhizal species, i.e., *A. thaliana* (Fiorilli *et al*., 2019; Wang *et al*., 2019). *In vitro*, this enzyme cleaves the apocarotenoid 3-OH-β-apo-10’-carotenal (C_27_) at the C13-C14 double bond, generating zaxinone, a C_18_-ketone (3-OH-β-apo-13-carotenone) that acts as a growth regulator, and an unstable C_9_-dialdehyde (Wang *et al*., 2019). Loss-of-function *zas* mutant showed a decreased zaxinone content in roots, reduced shoot, and root growth, and a higher SL level compared to wild-type rice plants (Wang *et al*., 2019). Phylogenetic analyses revealed that the rice genome encodes three *OsZAS* homologs, named *OsZAS1b, OsZAS1c*, and *OsZAS2* (Ablazov *et al*., 2023). Intriguingly, although *OsZAS2* is placed in a clade different from that of ZAS, it catalyzes the same reaction, and both enzymes contribute to zaxinone production in rice (Ablazov *et al*., 2023).

Apocarotenoids play several roles in plants, from regulating root and shoot developmental processes to coordinating plant responses to abiotic and biotic stress (Moreno *et al*., 2021). They are also emerging signaling molecules implicated in plant-microbe interactions, including the arbuscular mycorrhizal (AM) symbiosis (Fiorilli *et al*., 2019). The AM symbiosis is one of the most ancient and widespread associations, formed by approximately 70% of land plants (Wang and Qiu, 2006; Brundrett, 2009), including major crops, with soil fungi belonging to the Glomeromycotina group (Spatafora *et al*., 2016). In this symbiosis, the fungus facilitates the plant uptake of minerals, predominantly phosphorus (P) and nitrogen (N) (Smith *et al*., 2011), and the tolerance to biotic and abiotic stress (Pozo *et al*., 2010; Chen *et al*., 2018). Meanwhile, the plant provides the fungus with fixed organic carbon. The establishment of the AM symbiosis includes several steps, starting with partner recognition *via* diffusible molecules that activate the common symbiosis signaling pathway (MacLean *et al*., 2017) and trigger the development of fungus adhesion structures, called hyphopodia, on the root epidermis. These structures permit the fungus to enter the host root tissues and proliferate within cells or intracellularly (Bonfante and Requena, 2011; Nadal and Paszkowski, 2013). Finally, fungal hyphae invade the inner cortical layers, penetrate single cells and form highly branched tree-shaped hyphal structures, the arbuscules, where nutrient exchanges occur (Harrison, 2012; Gutjahr and Parniske, 2013). During these stages, the plant controls fungal expansion and symbiotic functions, by activating a series of cellular, metabolic, and physiological changes (Gutjahr, 2014; Carbonnel and Gutjahr, 2014). Among the environmental factors that regulate AM colonization, phosphate (Pi) availability is certainly one of the most crucial ones (Smith *et al*., 2011; Richardson *et al*., 2011). It has been recently shown that a complex gene network centered on the plant Pi starvation response actively supervises AM fungal development in roots, acting at the local and systemic level (Shi *et al*., 2021; Das *et al*., 2022). Pi starvation also induces SL biosynthesis and release (Yoneyama *et al*., 2007; Wang *et al*., 2017, 2022), while high Pi levels repress the expression of genes involved in the biosynthesis of carotenoids and SLs in root (Carbonnel and Gutjahr, 2014; Haider *et al*., 2023). SLs are the best-known plant molecules active in the pre-symbiotic interaction with AM fungi. In Pi-starved plants, SLs are produced by roots and exported to the rhizosphere, which directly stimulates AM fungal metabolism, gene expression, and hyphal branching, supporting the development of this symbiosis (Waters *et al*., 2017; Müller and Harrison, 2019). Notably, (Volpe *et al*.) recently showed that SL biosynthesis is stimulated by chito-oligosaccharides released by AM fungi.

Studies of the last decade highlighted that other apocarotenoid compounds are involved in the AM symbiosis (Fiorilli *et al*., 2019 and reference therein), including the plant hormone ABA that is known for coordinating plant’s response to biotic and abiotic stress factors (Felemban *et al*., 2019; Moreno *et al*., 2021). ABA has been reported to be involved in mycorrhizal colonization in different host plants, probably through synergistic and antagonistic interactions with other hormones (Herrera-Medina *et al*., 2007; Martín-Rodríguez *et al*., 2011; Charpentier *et al*., 2014). Specifically, a positive correlation between ABA levels and SL biosynthesis was observed, suggesting that ABA and SLs collaborate to influence the outcome of the symbiosis (López-Ráez *et al*., 2010). In contrast, ABA controls the normal development of arbuscules by inhibiting ethylene production (Martín-Rodríguez *et al*., 2011) and acts as an antagonist of gibberellins (GA) by down-regulating their biosynthesis and promoting their catabolism (Floss *et al*., 2013; Martín-Rodríguez *et al*., 2016).

Blumenols (C_13_) and mycorradicin (C_14_) are further apocarotenoids associated with AM symbiotic establishment and maintenance (Walter *et al*., 2007; Floß *et al*., 2008; Floss *et al*., 2008; Fiorilli *et al*., 2019) and described as a signature for AM symbiosis because of their being specifically accumulated in mycorrhizal plants (Walter *et al*., 2007; Hill *et al*., 2018; Moreno *et al*., 2021). Mycorradicins cause typical yellow/orange pigmentation of roots, which enabled their identification (Scannerini and Bonfante-Fasolo, 1977; Klingner *et al*., 1995; Floss *et al*., 2008). Blumenols are accumulated in roots and shoots of host plants in direct correlation with the fungal colonization rate (Klingner *et al*., 1995; Maier *et al*., 1997; Walter *et al*., 2000; Fester *et al*., 2002; Strack and Fester, 2006). Even if their biological role has not yet been clarified, blumenols have been proposed as foliar markers that allow rapid detection of AM symbiosis and screening of functional AM associations (Walter *et al*., 2010; Wang *et al*., 2018). Finally, zaxinone, a recently discovered apocarotenoid growth regulator (Wang *et al*., 2019; Ablazov *et al*., 2020) is also involved in AM symbiosis and acts as a component of a regulatory network that includes SLs, as demonstrated by that the impact of the rice gene encoding Zaxinone Synthase (*OsZAS*) on the extent of AM colonization SLs (Votta *et al*., 2022).

In the current study, we further explored the involvement of apocarotenoids in the AM symbiosis. For this purpose, we generated a qualitative and quantitative profile of apocarotenoids in roots and shoots of rice plants exposed to high/low Pi concentrations (+Pi and -Pi) and upon AM symbiosis in a time course experiment covering different stages of growth and AM development. We complemented the metabolic profiles by characterizing the expression pattern of *CCD* genes and took advantage of chemometric tools to get deeper insights in the biology of apocarotenoids during this plant-fungal symbiosis.

## Materials and methods

### Plant and fungal materials

Rice seeds of wild-type (WT) (cv. Nipponbare) were germinated in pots containing sand and incubated for 10 days in a growth chamber under 14 h light (23 °C)/10 h dark (21 °C). A set of plants (MYC) was inoculated with *Funneliformis mosseae* (BEG 12, MycAgroLab, France). The fungal inoculum (15%) was mixed with sterile quartz sand and used for colonization. A group of non-mycorrhizal plants (no-myc -Pi) was also set up. These two groups of plants (MYC and no-myc -Pi) were watered with a modified Long-Ashton (LA) solution containing 3.2 μM Na_2_HPO_4_·12H_2_O (low Pi) and grown in a growth chamber under 14 h light (24 °C)/10 h dark (20 °C) regime. Another group of no-myc WT plants was watered with a LA containing 500 μM Na_2_HPO_4_·12 H_2_O (+Pi) and grown in the same condition described above; these plants were considered the no-myc + Pi samples. Plants for the three different conditions (MYC, no-myc -Pi, no-myc +Pi) were collected at three time points: 7 days post-inoculation (dpi), 21 dpi, and 35 dpi. For the molecular and metabolites analyses, roots and shoots samples were harvested and immediately frozen in liquid nitrogen and stored at -80° C.

### Qualitative and quantitative profiling of plant apocarotenoids (APOs)

Following the method used by Mi *et al*. (2018), about 20 mg lyophilized root and shoot tissue powder was spiked with Internal Standards (IS) mixture (2 ng each standard) and extracted with 2 mL of methanol containing 0.1% butylated hydroxytoluene (BHT) in an ultrasound bath (Branson 3510 ultrasonic bath) for 15 min, followed by the centrifugation. The supernatant was collected, and the pellet was re-extracted with 2 mL of the same solvent. The two supernatants were then combined and dried under vacuum. The residue was re-dissolved in 150 μL of acetonitrile and filtered through a 0.22 mm filter for LC-MS analysis.

Analysis of apocarotenoids was performed on a Dionex Ultimate 3000 UHPLC system coupled with a Q-Orbitrap-MS (Q-Exactive plus MS, Thermo Scientific) with a heated electrospray ionization source. Chromatographic separation was carried out on an ACQUITY UPLC BEH C_18_ column (100 × 2.1mm, 1.7 μm) with a UPLC BEH C18 guard column (5 × 2.1mm, 1.7 mm) maintained at 35° C. UHPLC conditions including mobile phases and gradients were optimized based on the separation of APOs and the time needed for sample analysis. APO isomers were identified by MS/MS fragmentation.

The quantification of APOs was calculated as follows: Amount [target APO] = Area [target APO]/Area [spiked IS] x Amount [spiked IS]/ mg materials. The experiment was repeated twice with equivalent results.

### Gene expression analysis

Total RNA was extracted from WT rice roots using the Qiagen Plant RNeasy Kit according to the manufacturer’s instructions (Qiagen, Hilden; Germany). Following the producer’s directives, samples were treated with TURBO™ DNase (Thermofischer). The RNA samples were routinely checked for DNA contamination through PCR analysis. Single-strand cDNA was synthesized from 1 μg of total RNA using Super-Script II (Invitrogen) according to the instructions in the user manual. Quantitative RT-PCR (qRT-PCR) was performed using a Rotor-Gene Q 5plex HRM Platform (Qiagen). All reactions were performed on at least three biological and three technical replicates. Baseline range and take-off values were automatically calculated using Rotor-Gene Q 5plex software. The transcript level of genes listed in **Supplemental Table 1** was normalized using *OsRubQ1* housekeeping gene (Güimil *et al*., 2005). Only take-off values leading to a Ct mean with a standard deviation below 0.5 were considered. The experiment was repeated twice with equivalent results.

### Statistics and reproducibility

Both experiments (plant apocarotenoid quantification and CCD gene expression analysis) were performed with at least three biological replicates each. Statistical tests were carried out through One-way analysis of variance (One-way ANOVA) and Tukey’s *posthoc* test, using a probability level of P<0.05. All statistical elaborations were performed using PAST statistical package version 4 (Hammer *et al*. 2001).

### Data quality assessment and preprocessing

Gene and apocarotenoid datasets were inspected to spot potential extreme samples or outliers. Different preprocessing approaches were tested, including autoscaling (i.e., column scaling to unit variance, followed by mean centering) and normalization to a unit area (i.e., the normalization factor of each sample was computed from its “area under the curve”) followed by mean centering. Based on the ease of interpretation, we selected the following preprocessing: mean centering alone for the apocarotenoids dataset, and autoscale for the gene dataset. All modeling results were therefore obtained from the two datasets preprocessed as such. With respect to the analysis with the low-level data fusion approach (i.e., combining the two datasets into an individual one), a different sequence tailored to the issue of obtaining an equal representation of the two datasets was used, as described in the dedicated paragraph further in the section Data fusion approach.

### Exploratory analysis

All chemometric models reported in this work are “exploratory”, meaning that they describe the phenomena and natural groupings captured in the data, in an unsupervised manner (Li Vigni *et al*. 2013). To this aim, Principal Component Analysis (PCA) (Bro & K. Smilde 2014) was employed. This technique is employed to capture, in sequence, the largest sources of variability by defining new variables (the so-called “Principal Components”, PCs), which are summaries of the different pieces of information contained in the data. This “summarized” version of the information can be inspected with the scores and loadings plots, which are scatter plots obtained by plotting pairs (and sometimes triplets) of PCs. The scores plot allows inspecting the relationships among the samples and thus spotting possible groupings and tendencies or patterns of interest, while the loadings plot allows inspecting the relationships among the variables of the data, providing at the same time an interpretation of the scores plot. In our study, individual apocarotenoids and genes were identified by their systematic names and inspected in PCA as the samples. At the same time, the variables of the datasets were the combinations of three-time points (7, 21, and 35 dpi) and three conditions (MYC, no-myc -Pi, no-myc +Pi) for a total of nine combinations.

### Data fusion approach

For this study we also tested a low-level data fusion approach (Borràs *et al*. 2015) to combine and jointly explore the information of the apocarotenoids and genes datasets. In practice, the apocarotenoids dataset was joined with the gene dataset in the sample direction so that the nine variables (combinations of time points and conditions) were coherent between the two datasets, i.e., the information described by each column had to be the same in both datasets.

We performed the following data preprocessing and fusion sequence: (i) standard deviation scaling for each APO/gene quantification, (ii) fusion of the two data tables, (iii) group scale to give the two data tables the same importance (i.e., each dataset accounts for 50 % of the total variance of the resulting fused dataset), (iv) mean center. The new fused data table was then modelled with PCA, with the apocarotenoids and the genes as the samples (rows) and the combinations of time points and treatments as the variables (columns) (**Supplementary Fig.S1**).

## Results

### *CCD* gene expression pattern during AM symbiosis

We determined the transcript level of a set of *CCD* gene, including *CCD1, CCD4a, CCD4b, CCD7, CCD8, ZAS1, ZAS1b, ZAS1c*, and *ZAS2*, in mycorrhizal plants grown at low Pi (3.2 μM) and in non-mycorrhizal plants grown at low (3.2 μM) or high Pi (500 μM). We measured the transcript levels at early (7 dpi), middle (21 dpi), and late (35 dpi) stage of AM symbiosis development (**Supplementary Fig. S2**). To assess the statistically significant differences, all samples were referred to the -Pi condition within each time point. As shown in the heatmap (**Fig. 1A**) referred to roots, *CCD1, CCD4a*, and *CCD4b* transcript level increased at 21 dpi in the +Pi condition compared to -Pi and MYC ones. Concerning the SL biosynthetic genes, we observed an induction of *CCD7* in MYC roots at the middle and late stage (21, 35 dpi), while *CCD8* was induced at 7 dpi under the MYC condition, and, as expected, down-regulated under +Pi condition in the later time points (at 21 and 35 dpi) (López-Ráez *et al*., 2008; Yoneyama *et al*., 2013). At the middle stage (21dpi), *ZAS1* showed an up-regulation in MYC samples and a down-regulation under +Pi. The *ZAS1* homolog, *ZAS1b* was upregulated at 21 dpi in MYC and +Pi roots, while we detected a down-regulation in MYC roots during the later stage. By contrast, *ZAS1c* was up-regulated at 35 dpi in MYC roots. Finally, *ZAS2* expression level increased at 21 dpi in +Pi. **Supplementary Fig. S3A**.

**Figure 1.**
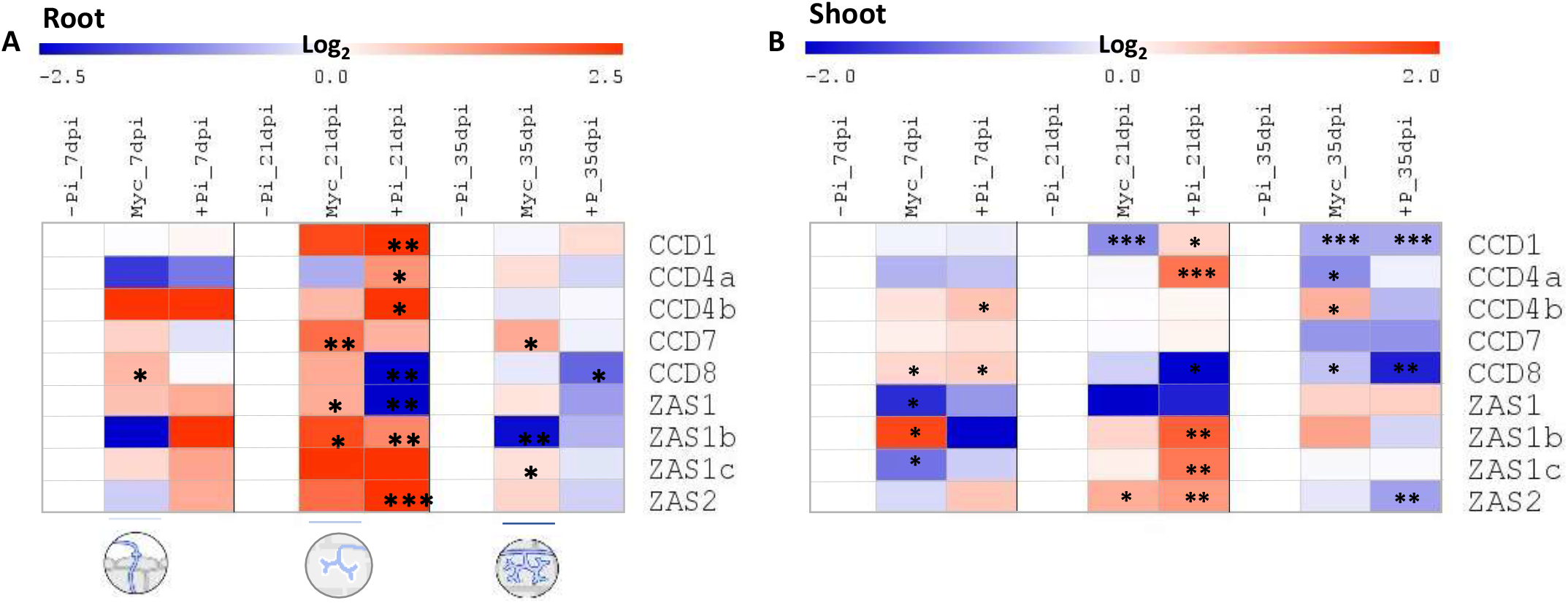
Heatmap of root (A) and shoot (B) gene expression of the three-time points (7, 21, 35 dpi: days post inoculation) and the three analyzed conditions (-Pi, MYC, +Pi). Data are means ± SE (n<=4). For each gene and time point, the value of the corresponding -Pi sample was set to 1. Asterisks indicate statistically significant differences referred to the –Pi condition, separately for each time point, by one way-Anova (*P < 0.05; **P<0.01; ***P<0.001). The circles represented the different stages of mycorrhization: at the early stage (7 dpi), the fungus structures, hyphopodia, adhered to the root epidermis, during the middle stage (21 dpi), the arbuscules started their development that will be completed at the later stage (35 dpi). Heatmaps were generated with the MultiExperiment Viewer (MeV) software. -Pi: 3.2 μM Pi; myc: mycorrhizal plants grown at 3.2 μM Pi; +Pi:500 μM Pi.

*CCDs* gene expression pattern in shoots (**Fig. 1B, Supplementary Fig. S3B**) displayed several differences compared to roots. We detected induction of *CCD1* at 21 dpi in the +Pi condition, while its expression decreased in the MYC condition at 21 and 35 dpi. *CCD4a* showed an expression profile similar to *CCD1*. By contrast, *CCD4b* displayed an opposite trend compared to *CCD4a* with an up-regulation at 35 dpi in the MYC condition. Moreover, *CCD4b* displayed an up-regulation at 7 dpi in the +Pi condition. We did not observe significant changes in the *CCD7* expression level across all conditions or time points, while *CCD8* transcript level was up-regulated at 7 dpi (MYC and +Pi conditions) and down-regulated in the later stages (21 and 35 dpi) in shoots of plants grown in +Pi and at 35 dpi in shoots of MYC plants. *ZAS1* displayed a down-regulation trend in all time points and conditions considered, with a statistically significant difference in mycorrhizal samples at 7 dpi. *ZAS1b*, and *ZAS1c*, showed an up-regulation in leaves of the MYC plant at the first time point (7 dpi) and 21 dpi upon +Pi. Lastly, *ZAS2* was barely detected in the shoot of plant growth at low Pi, while it showed an up-regulation at 21 dpi in MYC and +Pi conditions and a down-regulation in +Pi at 35 dpi.

We used PCA to assess the samples’ natural grouping and clustering tendencies under the different growth conditions (MYC, -Pi, and +P) at the three-time points analyzed. The loading plot of **Fig. 2A** describes the influence of the measured variables on the samples’ distribution shown in the scores plot of **Fig. 2B** that provides insights into this distribution.

**Figure 2.**
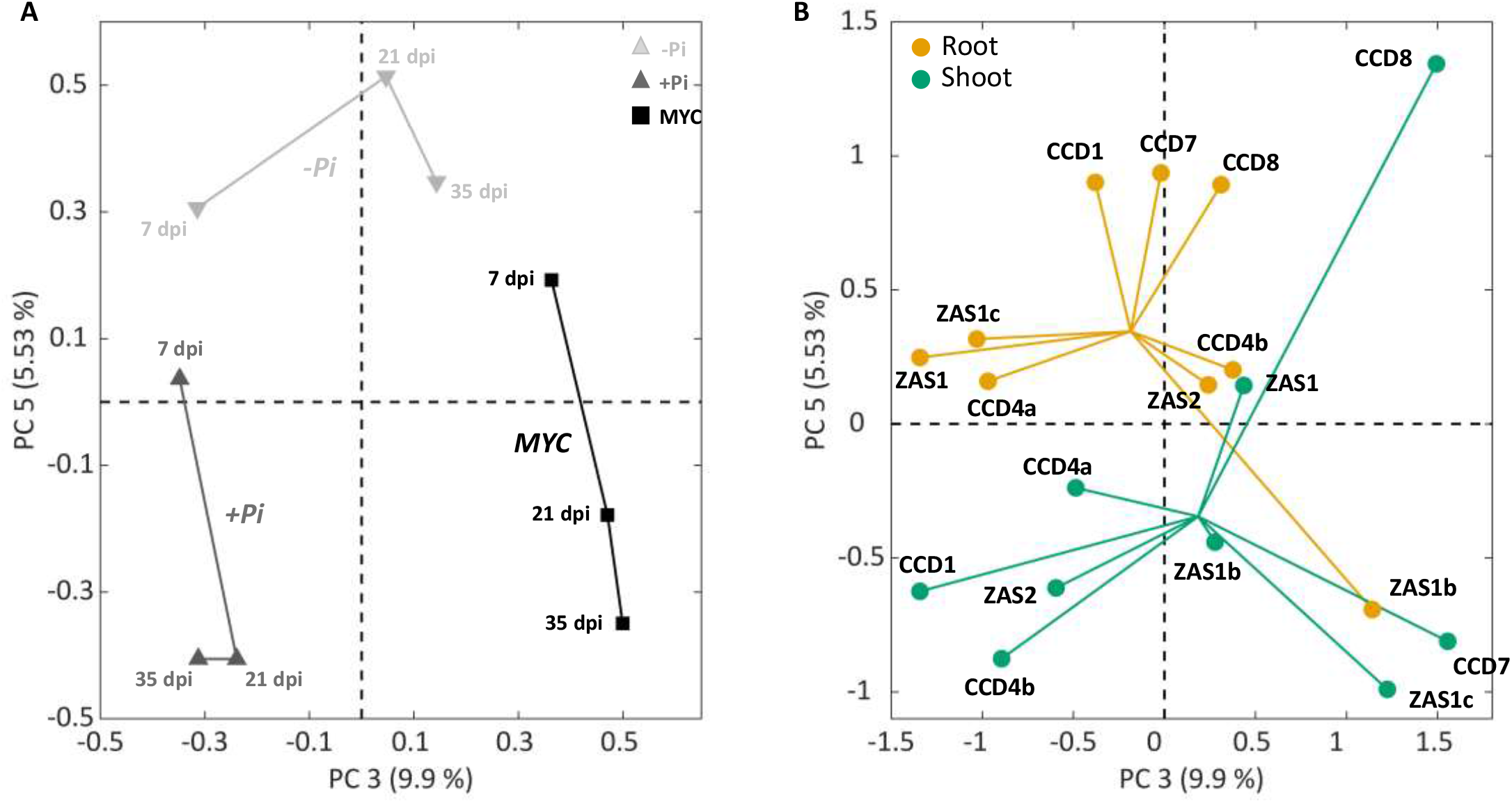
Principal component analysis of root and shoot CCDs genes expression across the three-time points and the three growth conditions (-Pi, MYC, +Pi). Loadings plot (A) and scores plot (B) show the third and fifth principal components. In the scores plot (B) for both groups the lines connecting each sample lead to the cluster center.

In our study, the PCA model was obtained from the expression level of the set of genes analyzed in all samples, in which PC1 explained 43.88% (related to the gene expression level) and PC2 explained 22.23% (related to the plant developmental stage) of the total variance (**Supplementary Fig. S4**). PC1 and PC2 models highlight that in all growth conditions the middle developmental stage (21 dpi) presented a different *CCD* expression pattern compared to early and late stages. We selected the model with 5 PCs, since PC3 (9.90%) and PC5 (5.53%) provided notable information related to the growth conditions, as shown in the loading plot (**Fig. 2A)**, while the scores plot (**Fig. 2B**) displayed a clear separation between the plant organs: roots and shoots. More in detail, **Fig. 2B** showed that *CCD1* and *CCD7* expression in roots was mainly influenced by Pi level, while that of *CCD8* was affected by both Pi level and MYC condition. *CCD4a, ZAS1c*, and *ZAS1* expression was more affected by the time point than by the Pi level. By contrast, the *CCD4b* and *ZAS2* expression levels were mainly influenced by the Pi level.

Concerning the shoot, *CCD1, CCD4a, CCD4b*, and *ZAS2* were located in the plot area influenced by the Pi level during the middle and late stages (21 and 35 dpi), while *CCD7* and *ZAS1c* fell in the plot area related to the MYC condition.

### Apocarotenoid profile

To profile non-hydroxylated and hydroxylated apocarotenoids (**Table 1**) in roots and shoots of rice plants grown in high/low Pi concentration (+Pi and -Pi) and upon MYC condition, we used the ultra-HPLC (UHPLC)-mass spectrometry (MS)-based approach to get insight into the apocarotenoid compositions (Mi *et al*., 2018). To simplify this analysis, the statistically significant differences were referred to as the -Pi condition within each time point. The results showed a substantial difference between roots and shoots in the apocarotenoid quantification and distribution, in analogy to what has been observed in *CCDs* gene expression data.

**Table 1.**
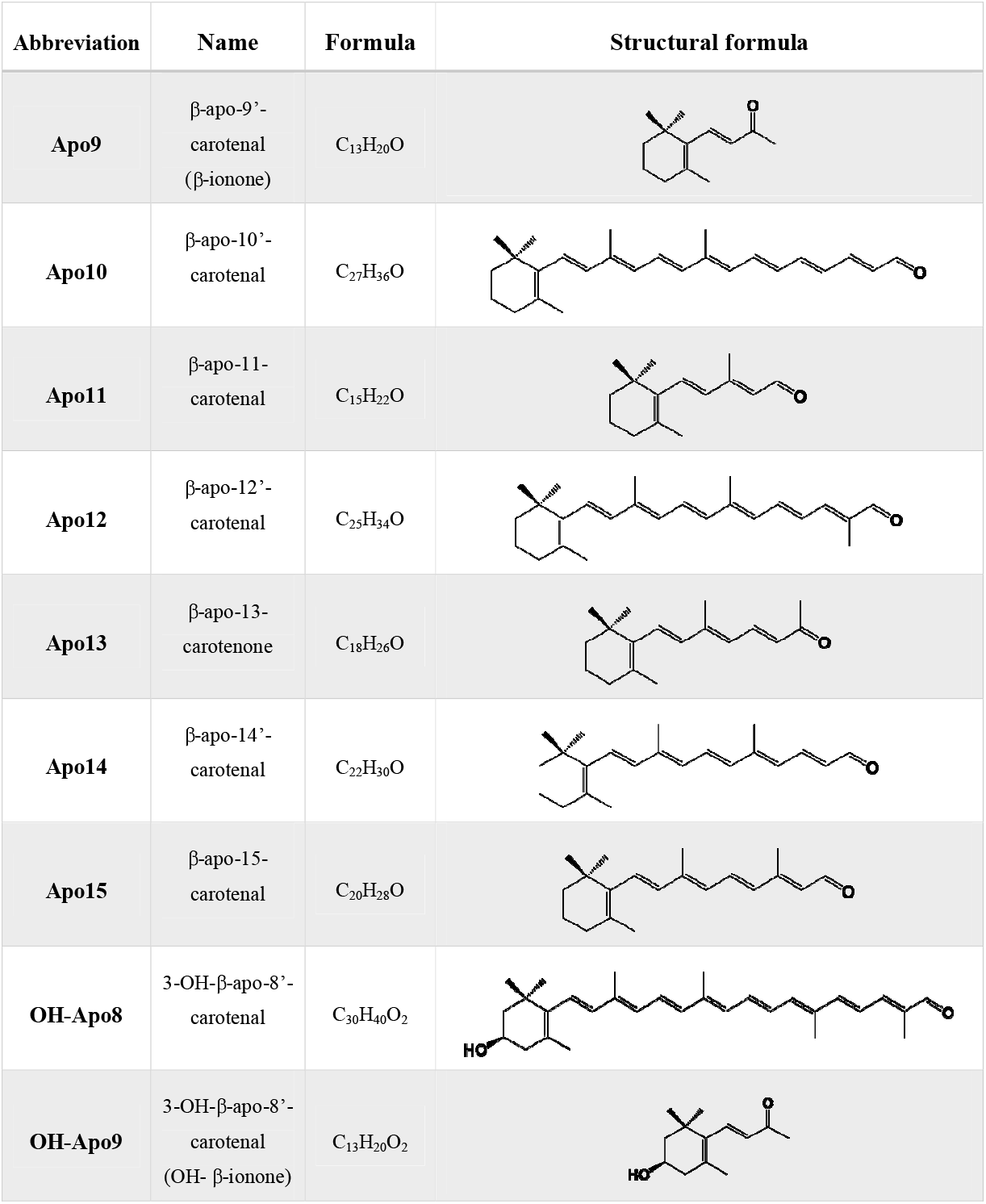
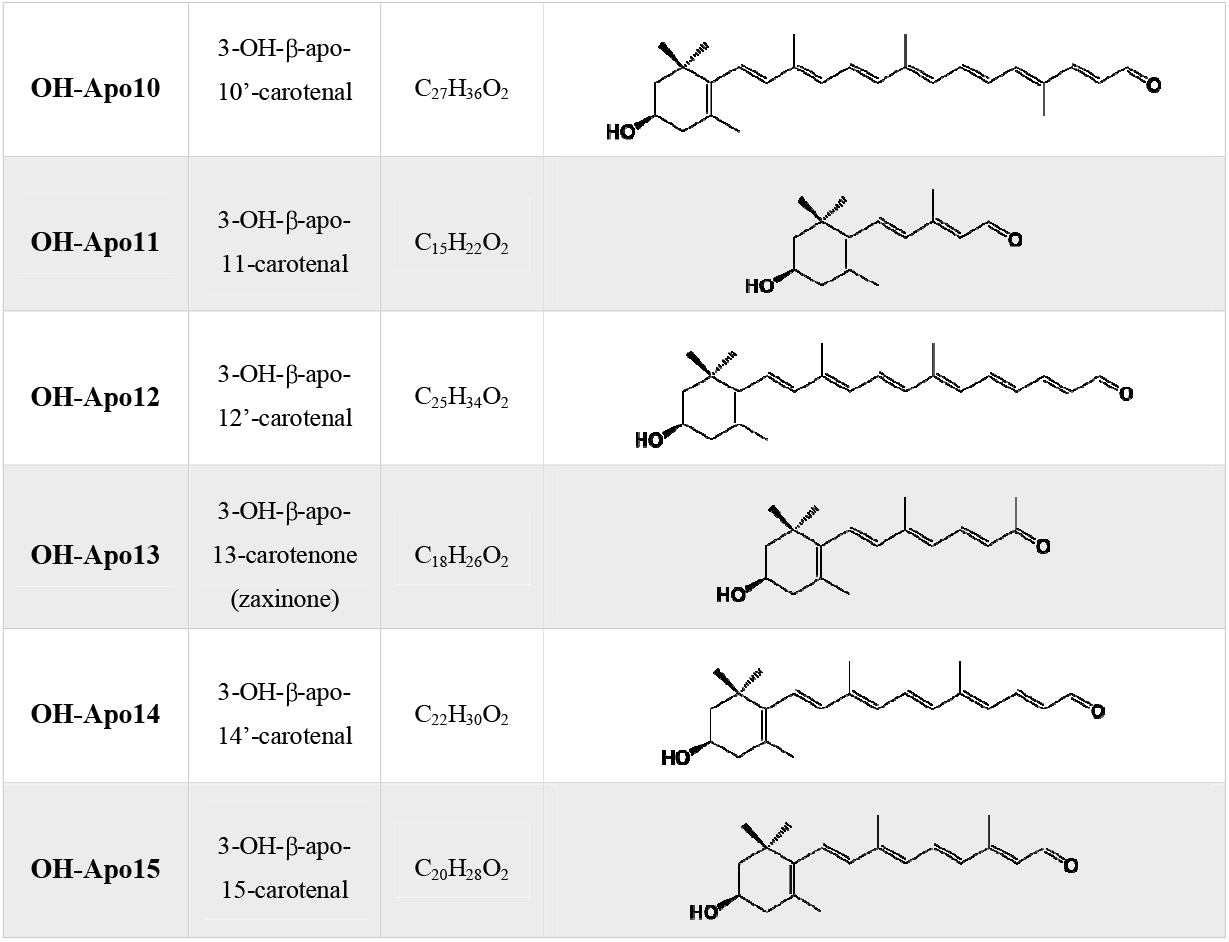
A summary with non-hydroxylated and hydroxylated apocarotenoids analyzed in this study, the formula, corresponding name, and structural formula are indicated for each abbreviation.

| Abbreviation | Name | Formula | Structural formula |
| --- | --- | --- | --- |
| **Apo9** | β-apo-9'-carotenal<br>(β-ionone) | $C_{13}H_{20}O$ | |
| **Apo10** | β-apo-10'-carotenal | $C_{27}H_{36}O$ | |
| **Apo11** | β-apo-11'-carotenal | $C_{15}H_{22}O$ | |
| **Apo12** | β-apo-12'-carotenal | $C_{25}H_{34}O$ | |
| **Apo13** | β-apo-13'-carotenone | $C_{18}H_{26}O$ | |
| **Apo14** | β-apo-14'-carotenal | $C_{22}H_{30}O$ | |
| **Apo15** | β-apo-15'-carotenal | $C_{20}H_{28}O$ | |
| **OH-Apo8** | 3-OH-β-apo-8'-carotenal | $C_{30}H_{40}O_2$ | |
| **OH-Apo9** | 3-OH-β-apo-8'-carotenal<br>(OH- β-ionone) | $C_{13}H_{20}O_2$ | |
| **OH-Apo10** | 3-OH-β-apo-10'-carotenal | $C_{27}H_{36}O_2$ | |
| **OH-Apo11** | 3-OH-β-apo-11-carotenal | $C_{15}H_{22}O_2$ | |
| **OH-Apo12** | 3-OH-β-apo-12'-carotenal | $C_{25}H_{34}O_2$ | |
| **OH-Apo13** | 3-OH-β-apo-13-carotenone<br>(zaxinone) | $C_{18}H_{26}O_2$ | |
| **OH-Apo14** | 3-OH-β-apo-14'-carotenal | $C_{22}H_{30}O_2$ | |
| **OH-Apo15** | 3-OH-β-apo-15-carotenal | $C_{20}H_{28}O_2$ | |

With respect to roots (**Fig. 3A, Supplementary Fig. S5A**), we observed an increment of the content of Apo9 (β-ionone), Apo10, and their hydroxylated forms (OH-Apo9 and OH-Apo10) in MYC condition at 21 dpi and 35 dpi. In addition, Apo10 showed a higher accumulation in MYC roots and upon +Pi at 7 dpi. Likewise, at the same time point, the Apo11 level increased in the MYC condition, while it decreased during the middle stage (21 dpi), contrary to its hydroxylated forms (OH-Apo 11 and OH-Apo 11-iso) that showed a strong accumulation. Moreover, at 35 dpi, all the β-apo-11-carotenoids (C_15_) showed a statistically significant decrease upon +Pi. Apo12 and Apo14 displayed the same pattern at 35 dpi: both showed a higher accumulation in MYC and +Pi compared to -Pi. By contrast, we observed an increase of Apo13 in MYC and +Pi conditions during the early (7 dpi) and middle stages (21 dpi); its hydroxylated forms (OH-Apo13 and OH-Apo13-iso) also displayed a higher content at 7 dpi in the +Pi condition and 21 dpi in the MYC root. Moreover, at the later stage (35 dpi), OH-Apo13-iso showed a statistically significant higher and lower content in MYC and +Pi roots respectively. Finally, Apo15 and Apo15-iso displayed a higher level at 21 dpi in MYC and +Pi roots.

**Figure 3.**
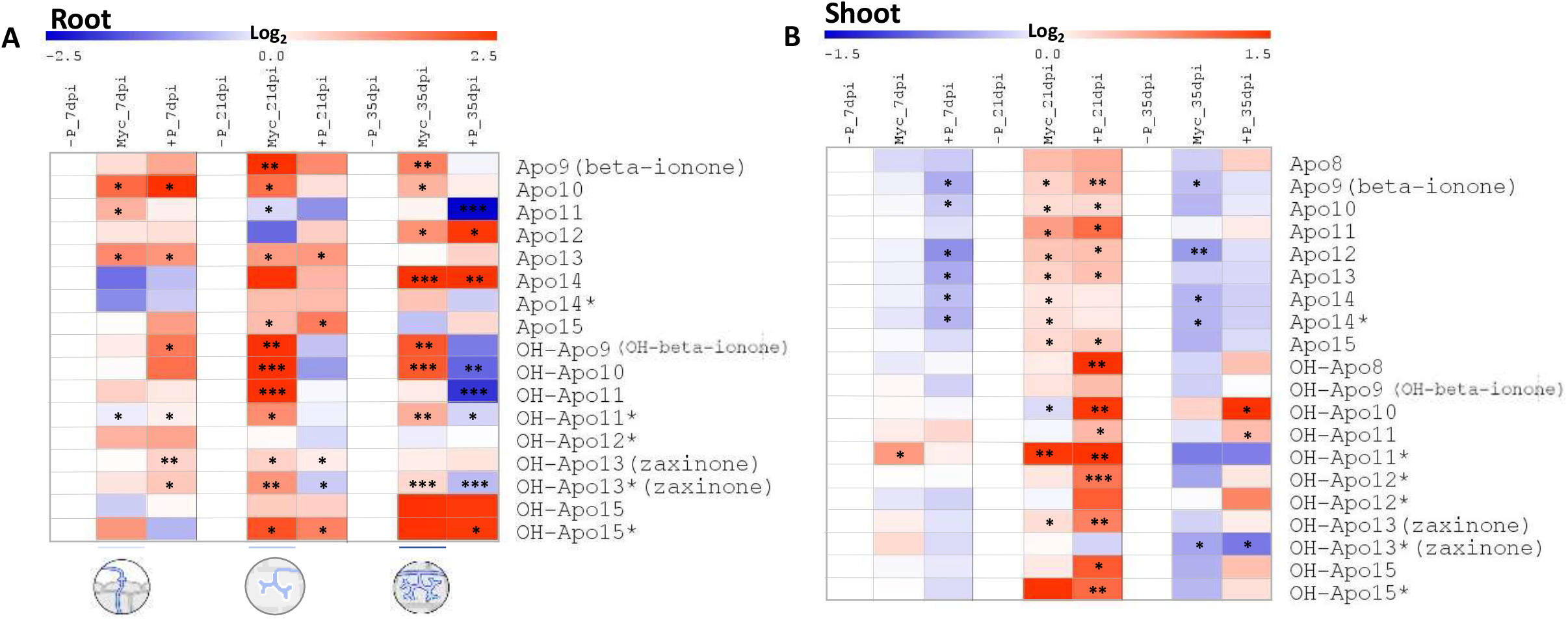
Heatmap of root (A) and shoot (B) apocarotenoids quantification across the three time points and the three analyzed conditions (-P, MYC, +P). For each APOs and time point, the value of the corresponding -Pi was set to 1. Data are means ± SE (n<=4). Asterisks indicate statistically significant differences as compared to –P condition, separately for each time point, by one way-Anova (*P < 0.05; **P<0.01; ***P<0.001). The circles represented the different stages of mycorrhization: at the early stage (7 dpi), the fungus structures, called hyphopodia, adhered to the root epidermis, during the middle stage (21 dpi), the arbuscules started their development that will be completed at the later stage (35 dpi). The apocarotenoids indicated with asterisks represented the isoform of the corresponding apocarotenoid. Heatmaps were generated with the MultiExperiment Viewer (MeV) software.

Concerning the shoot, the heatmap (**Fig. 3B**) showed that the non-hydroxylated apocarotenoids (from Apo 8 to Apo 15) displayed an overall similar profile: a general decrease at 7 and 35 dpi in +Pi condition compared to plants grown at -Pi, and an increase upon MYC and +Pi during the middle stage (21 dpi). Notably, Apo9, Apo10, Apo11, Apo12, Apo13, Apo14 and its isomer (Apo14-iso), and Apo15 levels decreased at 7 dpi in the +Pi condition. By contrast, at 21 dpi, Apo9, Apo10, Apo11, Apo12, Apo13, and Apo15 content increased in MYC and +Pi conditions, while Apo14 and its isomer showed an increment only for the MYC condition. At the later time point (35 dpi), we detected a decrease of Apo9, Apo12, and Apo14 e its isomer content in MYC plants.

The hydroxylated forms showed a profile similar to non-hydroxylated APO with an increased content at 21 dpi in the +Pi condition, and a decreasing trend in the MYC condition at 35 dpi. In detail, at 7 dpi, the OH-Apo11 isomer showed a statistically different increase in MYC condition. Further, at 21 dpi, OH-Apo8, OH-Apo10, OH-Apo11 and its isomer, OH-Apo12, OH-Apo13, OH-Apo15, and OH-Apo15 isomer strongly increase in +Pi. At the same time point, also OH-Apo11 isomer and OH-Apo13 levels increased in the MYC condition, while the OH-Apo10 content decreased. At the later stage, OH-Apo10 and OH-Apo11 displayed an increased content at +Pi, while OH-Apo13-iso accumulation decreased in the shoot of plants grown in MYC and +Pi conditions (**Supplementary Fig. S5B**).

To highlight correlations in apocarotenoids distribution across different stages of plant development, AM symbiosis and Pi levels, a PCA was employed. In the apocarotenoid database, no outliers (i.e., samples with clearly inconsistent values and/or unexpected behaviors attributable to errors of measurement or to data acquisition problems) were identified, even if three apocarotenoids (OH-Apo10, Apo10, and Apo12) showed very high values across all time points and treatments (**Supplementary Fig. S6**). To better model the information of the rest of the samples, these three extreme samples were removed from the apocarotenoid dataset and projected at a later stage to inspect their position in the final PCA model.

In the PCA model referred to apocarotenoids, PC1 explained 43.88% (related to apocarotenoid quantification) and PC2 explained 22.23% (related to growth conditions) of the total variance (**Supplementary Fig. S7**). PC1 and PC2 models highlighted the apocarotenoids strictly related to MYC condition (OH-Apo9; Apo11; OH-Apo11; OH-Apo13iso). Further, we adopted the PCA model PC2 combined with PC3, where PC3 explained 2.55% of the total variation, upon the propensity of samples to regroup following the temporal trend described in the loading plot (**Fig. 4A**).

**Figure 4.**
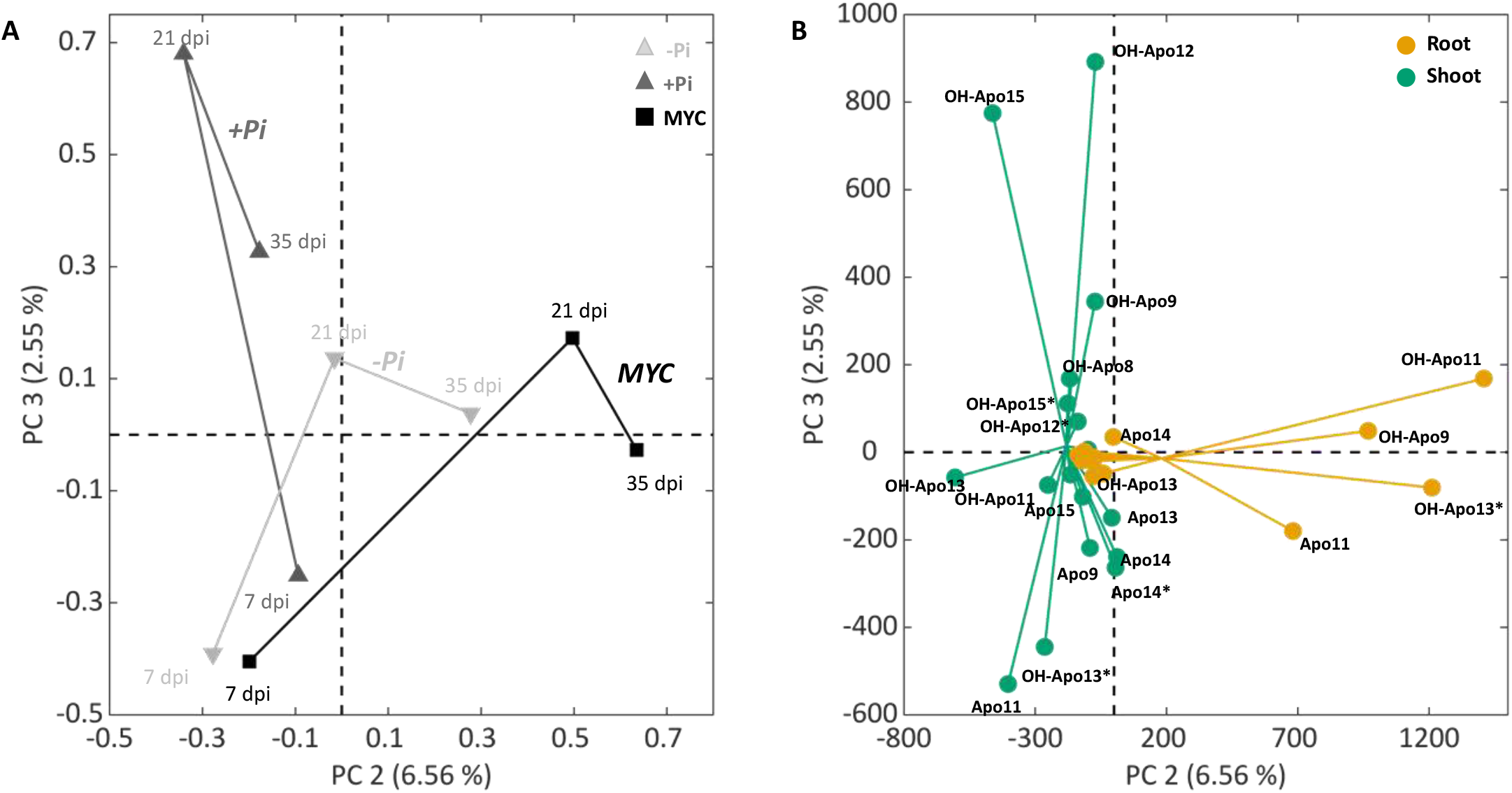
Principal component analysis of root and shoot APOs across the three-time points and the three growth conditions (-Pi, MYC, +Pi). Loadings plot (A) and scores plot (B) show the second and third principal components. In the scores plot (B) for both groups the lines connecting each sample lead to the cluster center.

In the plot chart, at 7 dpi all the growth conditions were clustered in the same area. Here, the majority of the analyzed shoot apocarotenoids (Apo9, Apo11, OH-Apo11 isomer, OH-Apo13, OH-Apo13 isomer, and Apo15) were located, suggesting that their content was mainly influenced by the growth time than by the growth condition.

The scores plot (**Fig. 4B)** displayed a clear separation between the apocarotenoids quantified in roots or shoots. In-depth, in root OH-Apo9, OH-Apo11, Apo11, and OH-Apo13 isomer were mainly influenced by mycorrhization in different time points (21 dpi and 35 dpi). Instead, APO14 seems to be dependent on the Pi level and MYC condition. However, in shoot OH-Apo9, OH-Apo12, and OH-Apo15 were linked to the +Pi condition. OH-Apo8, the OH-Apo12 isomer, and the OH-Apo15 isomer depended on the Pi level at the middle stage (21 dpi).

### Data fusion

Finally, to combine and investigate the potential correlation between apocarotenoids and CCD genes, we used a low-level data fusion approach (Borràs *et al*., 2015) to combine the two datasets into an individual fused one, also modelled with PCA.

In the resulting PCA model, considering genes expression and apocarotenoids profiles joined, PC1 explained 29.65% and PC2 explained 26.81% of the total variance (**Fig. 5A, Fig. 5B**). From the loading plot (**Fig. 5A**) we observed the grouping of the samples interpretable by the temporal trend (7, 21, and 35 dpi). As reported for the previous scores plots referred to individual categories (genes and apocarotenoids), in **Figure 5B** we observed a clear separation between plant organs. In more detail, genes and apocarotenoids in the left upper part of the scores plot (**Fig. 5B**) were more related to the early stage (7 dpi). Here, we found mainly shoot apocarotenoids, *CCD8* expressed in both root and shoot, and *ZAS1c* in the root. By contrast, the lower left part of the plot, clustered exclusively apocarotenoids and genes (*CCD7* and *ZAS1*) modulated in the root, depended on the MYC condition during the middle and later stages. In particular, Apo9 (β-ionone), Apo10, and their hydroxylated forms (OH-Apo9 and OH-Apo10) were grouped in this plot area, suggesting their possible involvement during the AM colonization process. Furthermore, in the same area, we highlighted the association between *CCD7* and one of its cleavage products, Apo9 (β-ionone). In addition, this group highlighted the correlation between *ZAS1*, responsible for the OH-Apo13 (zaxinone) synthesis, and its precursors (OH-Apo10 and OH-Apo12).

**Figure 5.**
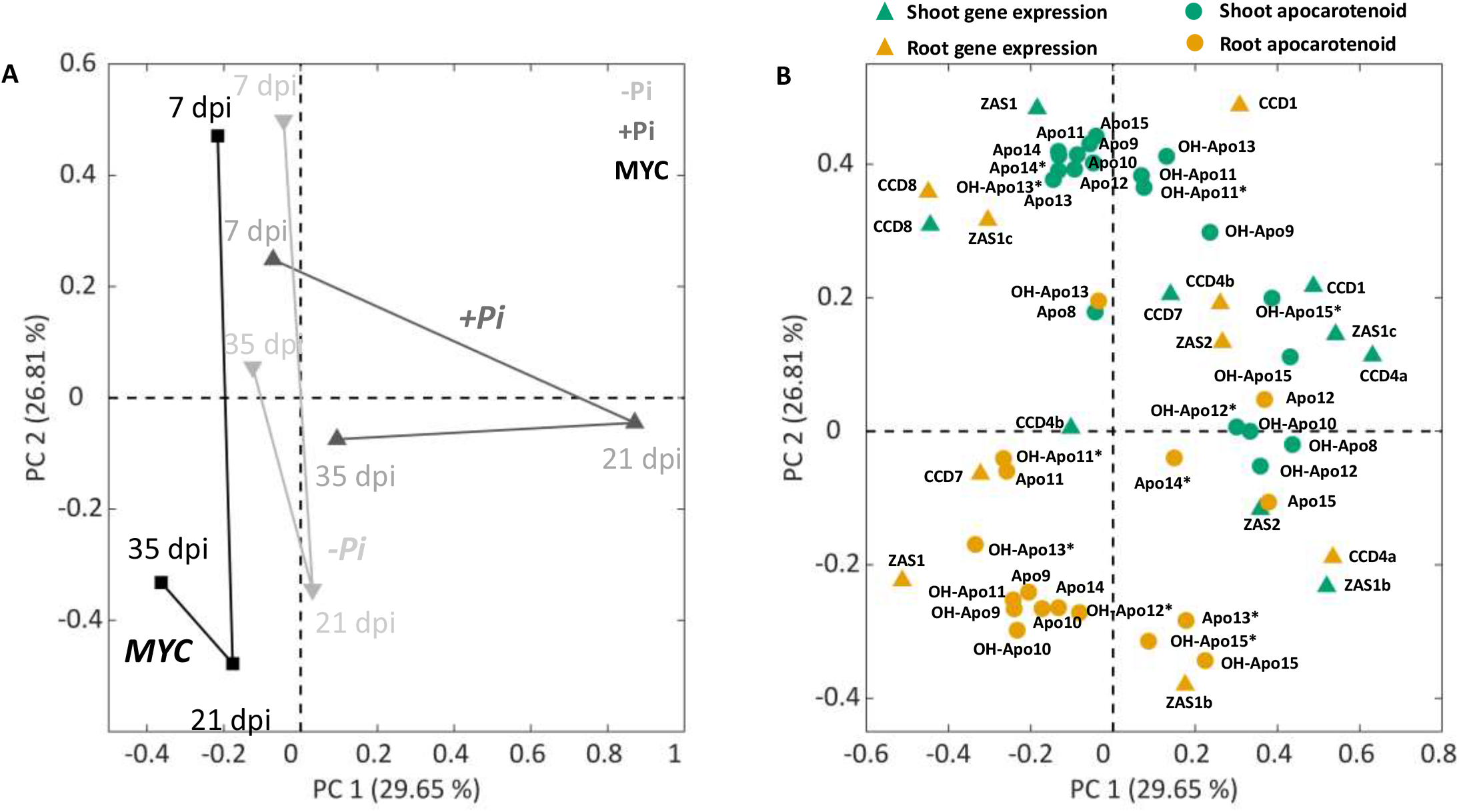
Principal component analysis of root and shoot genes and apocarotenoids datasets fused and analysed across the three-time points and the three growth conditions (-Pi, MYC, +Pi). Loadings plot (A) and scores plot (B) show the first and second principal components. The apocarotenoids indicated with asterisks represent the isoform of the corresponding apocarotenoid.

In the shoot, most of the *CCD* genes (*CCD1, CCD4a, CCD7, ZAS1b, ZAS1c*, and *ZAS2*) were mainly influenced by the Pi level and by the growing time (21 dpi and 35 dpi) similarly to some genes (*CCD1, CCD4a, CCD4b*, and *ZAS2*) in the root. Moreover, the right area of the plot clustered the majority of shoot apocarotenoids. On the whole, apocarotenoids profiles in the shoot seem to be more influenced by the time points (7, 21, 35 dpi) and Pi availability, while, in the root, most apocarotenoids and genes were mainly influenced by mycorrhization.

## Discussion

In recent years, plant apocarotenoids are emerging not only as carotenoid breakdown products but as metabolites with active roles in regulating physiological and developmental processes and plant-(a)biotic interactions (Zheng *et al*., 2021). In particular, some apocarotenoids were associated with the establishment and maintenance of AM symbiosis (Fiorilli *et al*., 2019). Investigations over the last decade indicate that beyond SLs, ABA, mycorradicins, and blumenols (Walter *et al*., 2007; Floss *et al*., 2008; Hill *et al*., 2018; Wang *et al*., 2018; Fiorilli *et al*., 2019), other apocarotenoids may play a role in this mutualistic association. For example, zaxinone, generated by the activity of the CCD subfamily Zaxinone Synthases, was shown to control the extent of AM root colonization with a complex interplay with SLs (Votta *et al*., 2022; Ablazov *et al*., 2023). To further explore the involvement of other apocarotenoids in the AM symbiosis in this work we developed a combined approach: we profiled apocarotenoids in rice roots and shoots across a time course (7, 21, and 35 dpi) experiment of AM colonization by LC-MS (Mi *et al*., 2018) and, in parallel, we monitored the expression pattern of a set of CCD genes. To highlight genes and apocarotenoids more specifically related to the AM association, we analyzed plants grown in low and high Pi conditions. Our results show that the AM colonization, although confined to the root system, can trigger a systemic response which is evident from the modulation of *CCDs* gene expression and apocarotenoid content in rice shoots. The effect on epigeous organs exerted by AM root colonization has been already described in other species (Fiorilli *et al*., 2009-2018; Zouari *et al*., 2014). In addition, our analysis indicates that both mycorrhization and Pi availability triggered an organ-specific response with differential modulation of genes and apocarotenoids in roots *versus* shoots (**Fig. 3, Fig. 4**).

In particular, in roots Apo9, OH-Apo9, Apo10 and OH-Apo10 are much more abundant in the MYC samples across almost all the time points analyzed and do not accumulate under +Pi, which suggests that they may be important for the AM colonization process. Apo9 is β-ionone, a cleavage product of several CCD enzymes (CCD1, CCD4, and CCD7) which was shown to have a role in plant-fungal interactions (Wilson *et al*., 1981; Sharma *et al*., 2012). However, its specific involvement in the AM symbiosis has not been characterized yet. As in our dataset, CCD7 displays an AM-responsive expression profile and it is involved in SLs biosynthesis, we envisage that β-ionone accumulation in AM roots is mainly due to CCD7 activity. It is worth noting that CCD7 could also be involved in the synthesis of blumenol-type metabolites and mycorradicin that accumulate in roots in the late stage of AM colonization (Wang *et al*., 2018; Fiorilli *et al*., 2019); we can also hypothesize that β-ionone is a precursor of blumenols. However, due to lack of authentic standards, we were not able to monitor blumenol derivatives.

Interestingly, we observed the accumulation at the middle and late stages of mycorrhization of zaxinone and OH-Apo10, which is the precursor of zaxinone, indicating that mycorrhization stimulates multiple steps of this branch of the apocarotenoid biochemical pathway. Notably, the expression of *ZAS1* (and partially *ZAS2*) was also highly influenced by the MYC condition in roots, confirming its correlation with zaxinone (**Fig. 1, Fig. 5**). The association between *CCD8* and *ZAS1* at 7 dpi is in line with previous data showing their interplay at the early stage of the AM symbiosis (Votta *et al*., 2022).

Apo11 level increased at 7 dpi and decreased at 21 dpi, while OH-Apo11 and OH-Apo11 isomers could be associated with the 21 dpi MYC condition. Importantly, the apocarotenoids Apo11 and OH-Apo11 were recently described as being part of an alternative zeaxanthin epoxidase-independent pathway to produce ABA (Jia *et al*., 2022). Moreover, these compounds act like ABA in maintaining seed dormancy and inducing the expression of ABA-responsive genes (Zheng *et al*., 2021; Jia *et al*., 2022). In light of these findings, we could hypothesize that Apo11 and OH-Apo11, could be involved in the AM symbiosis, and deserve more investigations along the whole colonization process and in relation to what has been already described for ABA in mycorrhizal roots (López-Ráez *et al*., 2010; Pozo *et al*., 2015).

The composition of shoot apocarotenoids seems to be most influenced by the time point considered. At 7 dpi, we observe a trend to a decrease of most apocarotenoids, especially in the +Pi compared to -Pi condition. At 21 dpi, the MYC and +Pi conditions showed a similar pattern with a general increase in the level of several apocarotenoids. At 35 dpi, MYC and +Pi conditions again displayed a similar profile, but with a general decrease of several apocarotenoids content. From these observations, it can be speculated that the similarity between the MYC and the +Pi conditions could mirror the Pi nutritional status since the AM symbiosis guarantees an improved Pi mineral nutrition mimicking the high Pi condition (Zouari *et al*., 2014). Moreover, our data indicate that the OH-Apo11 isomer deserves more investigation as it could be considered a shoot marker of the early stage of AM colonization.

We also attempted to associate the expression of specific genes with the accumulation of specific apocarotenoids with a data fusion approach as the genes involved in the production of many apocarotenoids are largely unknown. The reliability of the approach was confirmed by the association between *ZAS1* and zaxinone and its precursor in roots and by the correlation between *CCD7* and β-ionone (**Fig. 5**). In this context, we can speculate that *CCD1* and *CCD4a* are linked to the production of OH-Apo15 and its isomer, since they are correlated in the shoot and both organs, respectively (**Fig. 5**). Interestingly, a fungal *CCD, NosACO*, mediates Apo15 (retinal) production (Scherzinger *et al*., 2006), indicating that CCD1/4, or still unidentified CCDs, are involved in Apo15 formation during the AM symbiosis. In addition, the expression of *ZAS1* in shoots is also related to the accumulation of zaxinone, suggesting a direct involvement of this enzyme in endogenous zaxinone level in shoots.

In conclusion, our data show the specific profiles of *CCD* genes and apocarotenoids across different stages of the AM symbiosis and Pi conditions, possibly highlighting novel markers at both local and systemic levels. Moreover, this combined approach is a promising tool to further dissect this complex metabolic pathway, suggesting putative links between enzymatic activities and apocarotenoid production.

## Supplementary data

*Table S1*. List of primers used in this study.

*Fig. S1*. Data-fusion setup.

*Fig. S2*. Mycorrhization level in rice mycorrhizal plants across a time course (7, 21, and 35 dpi).

*Fig. S3*. qRT-PCR analysis of transcript levels of CCDs genes in rice root and shoot.

*Fig. S4*. Principal component analysis (PC1/PC2) of root and shoot CCDs genes across the three growth stages and conditions used in this study.

*Fig. S5*. Apocarotenoids quantification across the three time points and the three analyzed conditions.

*Fig. S6*. Plot of the raw apocarotenoids dataset, including the extreme values.

*Fig. S7*. Principal component analysis (PC1/PC2) of root and shoot apocarotenoids across the three growth stages and conditions used in this study.

## Acknowledgements

This work was supported by Baseline Funding and Competitive Research Grant (CRG2020) from King Abdullah University of Science and Technology given to SA-B. The authors would like to thank Jorge Gomez-Ariza for sharing the drawing of fugal structures present in Figure 1 and Figure 3.

## Author contributions

VF, SA-B, and LL designed and coordinated the investigation. CV and JYW performed the gene expression profiles and carried out the quantification of apocarotenoids with the help from KIL. NC and FS performed the PCA analysis. All authors contributed to the results and discussion. CV, VF, SA-B and LL wrote the article and all the authors contributed to manuscript review & editing.

## Data availability

The data supporting the findings of this study are available within the paper and within its supplementary data published online.

## Figure legends

**Fig. 1. Heatmap of root (A) and shoot (B) gene expression of the three-time points (7, 21, 35 dpi: days post inoculation) and the three analyzed conditions (-Pi, MYC, +Pi)**. Data are means

± SE (n<=4). For each gene and time point, the value of the corresponding -Pi sample was set to 1. Asterisks indicate statistically significant differences referred to the –Pi condition, separately for each time point, by one way-Anova (*P < 0.05; **P<0.01; ***P<0.001). The circles represented the different stages of mycorrhization: at the early stage (7 dpi), the fungus structures, hyphopodia, adhered to the root epidermis, during the middle stage (21 dpi), the arbuscules started their development that will be completed at the later stage (35 dpi). Heatmaps were generated with the MultiExperiment Viewer (MeV) software. -Pi: 3.2 μM Pi; myc: mycorrhizal plants grown at 3.2 μM Pi; +Pi: 500 μM Pi.

**Fig. 2. Principal component analysis of root and shoot CCDs genes expression across the three-time points and the three growth conditions (-P, MYC, +P)**. Loading plot (A) and scores plot (B) with the third and fifth principal components. In the scores plot (B) for both groups the lines connecting each sample lead to the cluster center.

**Fig. 3. Heatmap of root (A) and shoot (B) apocarotenoids quantification across the three time points and the three analyzed conditions (-P, MYC, +P)**. For each APOs and time point, the value of the corresponding -Pi was set to 1. Data are means ± SE (n<=4). Asterisks indicate statistically significant differences as compared to –P condition, separately for each time point, by one way-Anova (*P < 0.05; **P<0.01; ***P<0.001). The circles represented the different stages of mycorrhization: at the early stage (7 dpi), the fungus structures, called hyphopodia, adhered to the root epidermis, during the middle stage (21 dpi), the arbuscules started their development that will be completed at the later stage (35 dpi). The apocarotenoids indicated with asterisks represented the isoform of the corresponding apocarotenoid. Heatmaps were generated with the MultiExperiment Viewer (MeV) software.

**Fig. 4. Principal component analysis of root and shoot APOs across the three growth stages and conditions (-P, MYC, +P)**. Loading plot (A) and scores plot (B) with the second and third principal components. In the scores plot (B) for both groups the lines connecting each sample lead to the cluster center.

**Figure 5. Principal component analysis of root and shoot genes and apocarotenoids datasets fused and analyzed across the three time points and growth conditions (-P, MYC, +P)**. Loading plot (A) and scores plot (B) with the first and second principal components.

